# Compact attractors of an antithetic integral feedback system have a simple structure

**DOI:** 10.1101/868000

**Authors:** Michael Margaliot, Eduardo D. Sontag

**Author notes:** Research supported in part by research grants from the Israel Science Foundation, the US-Israel Binational Science Foundation, DARPA FA8650-18-1-7800, NSF 1817936, and AFOSR FA9550-14-1-0060.

## Abstract

Since its introduction by Briat, Gupta and Khammash, the antithetic feedback controller design has attracted considerable attention in both theoretical and experimental systems biology. The case in which the plant is a two-dimensional linear system (making the closed-loop system a nonlinear four-dimensional system) has been analyzed in much detail. This system has a unique equilibrium but, depending on parameters, it may exhibit periodic orbits. An interesting open question is whether other dynamical behaviors, such as chaotic attractors, might be possible for some parameter choices. This note shows that, for any parameter choices, every bounded trajectory satisfies a Poincaré-Bendixson property. The analysis is based on the recently introduced notion of *k*-cooperative dynamical systems. It is shown that the model is a strongly 2-cooperative system, implying that the dynamics in the omega-limit set of any precompact solution is conjugate to the dynamics in a compact invariant subset of a two-dimensional Lipschitz dynamical system, thus precluding chaotic and other strange attractors.

## I. INTRODUCTION

Feedback control is used by both natural and engineered systems, and in particular biological ones, in order to maintain the values of critical variables at precise homeostatic levels, or to force these variables to track reference signals so as to meet specifications. Examples of such variables at the cellular level include concentrations of proteins and other chemical species; at an organism level, variables such as blood sugar, blood pressure, water content, or temperature are often finely regulated. Feedback attenuates the degradation of performance due to disturbances as well as uncertainties not accounted for when originally designing (or evolving) a controller. Integral feedback (and more generally an “internal model” of the possible “exosignals) is necessary for regulation, as is well known in linear systems theory as well as for certain classes of nonlinear systems, and plays a central role in biological processes, both at the organism and cellular level [12]. Recently, there has been a considerable effort directed at the construction of integral feedback controllers in synthetic biology. One particular approach that has gained much recent attention, and the focus of this work, relies on two-molecule sequestration or annihilation reactions, and was introduced under the name antithetic controllers by Briat, Gupta, and Khammash [6]. Interestingly, the paper [4] shows mathematically that antithetic controllers are in a certain sense minimal-complexity controllers that achieve robust perfect adaptation.

Several groups have produced implementations of the antithetic architecture. In one of them, an *in vivo* construct was reported in [4] that is based on *Bacillus subtilis* sigW and its anti-factor rsiW as the two antithetic species. In another, in [2] we constructed and analyzed an *in vitro* synthetic biomolecular integral controller that precisely controls the protein production rate of an output gene, based on a sequestration reaction between *E. coli σ*_28_ and anti-*σ*_28_ factors.

Thus motivated, the purpose of this work is to analyze the following four-dimensional nonlinear dynamical system from [6]:

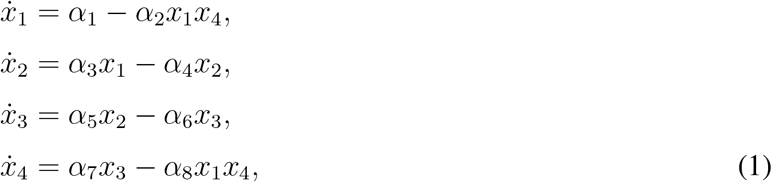

with all the *α*_*i*_’s being positive parameters. (In [6], *α*_2_ = *α*_8_, but we do not need that for the general theory.) We study this system in the nonnegative orthant 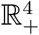, which is a (positively) invariant set of the dynamics and is the natural biological space to consider since these variables represent chemical species concentrations. This system is (with minor variations) also studied in [2], [1], where local stability and some partial global stability results are obtained.

The system (1) represents the closed-loop interconnection of an antithetic controller (described by the variables *x*_1_ and *x*_4_) and a simple two-dimensional linear system (described by the variables *x*_2_ and *x*_3_). It is easy to verify that this system has a unique equilibrium, for any (positive) parameter values. However, as shown in [6], a Hopf bifurcation may arise so that, depending on the parameters, the system can exhibit periodic orbits. An interesting open question is whether other dynamical behaviors, such as chaotic attractors, might be possible for some parameter choices. Our objective is to show that, *for any parameter choices*, every bounded trajectory satisfies a Poincaré-Bendixson property, thus precluding chaotic and other strange attractors. More generally, the same result holds for any “strongly 2-cooperative system”, of which the above system is a particular example, showing that the dynamics in the omega-limit set of any precompact solution is conjugate to the dynamics in a compact invariant subset of a two-dimensional Lipschitz dynamical system.

Here, we use the recently developed theory of *k*-cooperative dynamical systems [29] to analyze (1). Let 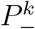 denote the set of vectors with no more than *k* − 1 sign variations (see the exact definition below). A linear time-varying dynamical system is called *k*-positive if its flow maps 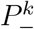 to 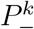, and strongly *k*-positive if its flow maps 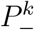 to the interior of 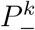. For *k* = 1, this reduces to the definition of a positive and strongly positive linear system [9]. A nonlinear system is called [strongly] *k*-cooperative if along any trajectory its associated variational system is [strongly] *k*-positive. For *k* = 1, this reduces to a [strongly] cooperative system [27].

The analysis of *k*-cooperative dynamical systems is based on the work of Sanchez [23] on dynamical systems with invariant cones of rank *k* (see also the more recent work [10]). A set *C* ⊂ ℝ^*n*^ is called a cone of rank *k* if: (1) *C* is closed, (2) *x* ∈ *C* implies that *ax* ∈ *C* for all *a* ∈ ℝ, and (3) *C* contains a subspace of dimension *k* and no subspace of a higher dimension. For example, it is straightforward to verify that 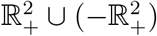 is an invariant cone of rank 1.

Sanchez [23] studied dynamical systems for which the flow of the associated variational equation maps an invariant cone of rank *k* to its interior. He showed that for some bounded trajectories the omega limit set can be projected in a one-to-one way to the invariant set of a Lipschitz dynamical system evolving on a *k*-dimensional space. For *k* = 2 this implies that these trajectories satisfy a Poincaré-Bendixson property. Roughly speaking, this means that any bounded trajectory that does not converge to an equilibrium converges to a limit cycle or a connection between equilibria.

It is not difficult to show that 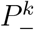 is an invariant cone of rank *k*. Thus, *k*-cooperative dynamical systems can be analyzed using the theory developed by Sanchez. However, the special structure of *k*-cooperative systems allows to deduce much more. Roughly speaking, the omega limit set of *every* bounded trajectory can be projected in a one-to-one way to the invariant set of a Lipschitz dynamical system evolving on a *k*-dimensional space.

Two important advantage of *k*-cooperative systems are: (a) the projection can be described in a simple and explicit form using the eigenvectors of a suitable matrix; and (b) the condition for *k*-cooperativity is an easy to check sign-pattern condition on the Jacobian of the vector field.

We show that the biological model (1) is a strongly 2-cooperative dynamical system, for any set of feasible parameter values. This then implies that its compact attractors consist of either equilibria, possibly connected by heteroclinic or homoclinic orbits, or closed orbits.

Some related work includes the following. Linear mappings that preserve the number of sign variations in a vector have been studied for a long time in the context of the variation diminishing property of totally positive (TP) matrices, that is, matrices with all minors positive [11]. Schwarz [24] extended these ideas to dynamical linear systems. We call a matrix *A* ∈ ℝ^*n*×*n*^ Jacobi if *A* is tridiagonal with positive entries on the super- and sub-diagonals. Schwarz [24] proved that the transition matrix of 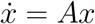, that is, exp(*At*) is TP for all *t* > 0 if and only if *A* is Jacobi. Since *x*(*t*) = exp(*A*(*t* − *t*_0_))*x*(*t*_0_) for all *t* ≥ *t*_0_ this implies that if *A* is Jacobi then the number of sign variations in *x*(*t*) can only decrease with *t*.

In a seemingly different line of research, Smillie [25] studied the nonlinear dynamical system 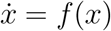 using the number of sign variations in the vector of derivatives 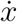 as an integer-valued Lyapunov function. He assumed that the Jacobian 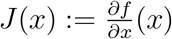 is a Jacobi matrix for all *x*. This was further developed by Smith [26] who considered time-varying and periodic nonlinear systems of a similar form. The recent paper [16] pointed out the connection between this work and the notion of a totally positive differential system (TPDS) introduced by Schwarz.

Mallet-Paret and Smith [15] studied a nonlinear system 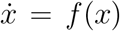 with a cyclic feedback structure, i.e. 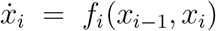, for *i* = 1, …, *n*, with *x*_0_ interpreted as *x*_*n*_. They showed that under certain assumptions on the signs of the nonzero entries of the Jacobian the number of *cyclic* sign variations in 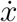 is a Lyapunov function and used this to prove a Poincaré-Bendixson property. Ref. [20] describes an algorithm for mapping measurements of state-variables to a dynamical system with a cyclic feedback structure. The theory of linear dynamical systems that preserve the number of cyclic sign variations, called *cyclic totally positive differential systems* (CTPDSs), was further developed in [5]. Elkhader [7] generalized the results of Mallet-Paret and Smith to a system in the form:

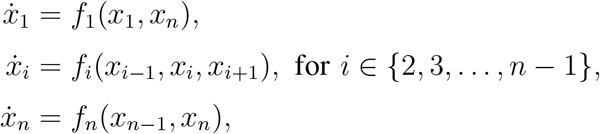

the dynamics for 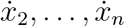 admits a tridiagonal structure (but not for 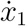), and there is a feedback connection from *x*_*n*_ to 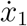 (but not from *x*_1_ to 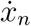). Again, he assumed certain sign conditions on the entries of the Jacobian. However, the biological model studied here admits a more general structure and it seems that it cannot be analyzed using the results above.

As noted in [13], [23], if *P* is a symmetric matrix with *k* negative eigenvalues and *n* − *k* positive eigenvalues then the set

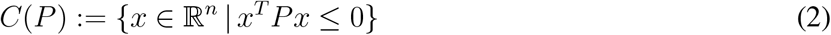

is a cone of rank *k*. This has been used to prove a Poincaré-Bendixson property for a certain type of dynamical systems in [28]. A recent paper [19] studied the same biological model studied as we do, using a a similar technique, but only for certain specific parameter values. Their analysis suggests that the model does not have a fixed *C*(*P*) as an invariant set. Rather, for different parameter values it admits different invariant sets in the form (2).

The remainder of this paper is organized as follows. The next section describes the general model that we consider, and provides our main result. The proof of this result is based on several auxiliary results given in Section IV. The main tool is the recently introduced notion of *k*-cooperative systems, and for the sake of completeness we first provide in Section III an introduction to the main definitions and results. The final section concludes and describes several directions for further research.

## II. THE MODEL

Consider the following nonlinear system:

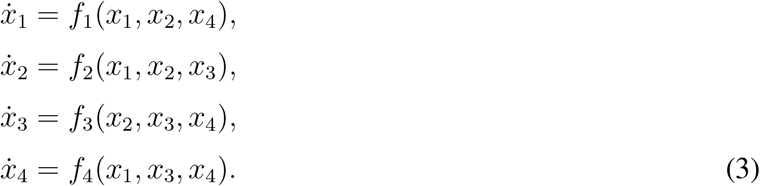

This represents a tridiagonal structure (i.e. every 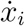 depends on *x*_*i*−1_, *x*_*i*_ and *x*_*i*+1_) plus a feedback connection from *x*_4_ to 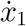 and from *x*_1_ to 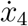. The state-variables describe quantities that can only take non-negative values. We assume that for any initial condition 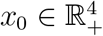 the corresponding solution *x*(*t, x*_0_) is unique and satisfies 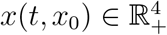 for all *t* ≥ 0.

The Jacobian 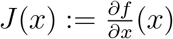 of the system is:

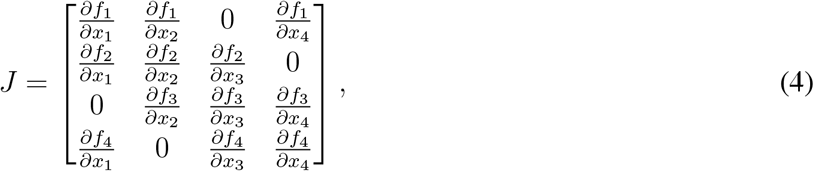

We pose several assumptions. First, for any 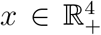 the Jacobian *J* (*x*) satisfies the following sign pattern:

a. 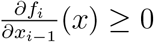 for *i* = 2,3,4;
b. 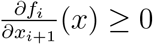 for *i* = 1,2,3;
c. 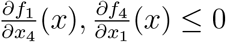

This implies that in the special case where *f*_1_ does not depend on *x*_4_ and *f*_4_ does not depend on *x*_1_ the system is cooperative, but in general the feedback from *x*_1_ to 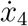 and from *x*_4_ to 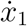 can be negative. Thus, *J* (*x*) is “almost” tridiagonal, with nonnegative entries on the super- and sub-diagonals, but it also has nonpositive values at entries (1, 4) and (4, 1). Note that *J* (*x*) is in general not a Metzler matrix. As we will see below, the assumption on the sign pattern of *J* implies that the system (3) is 2-cooperative.

Our second assumption is that for any 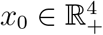 such that *x*(*t, x*_0_) remains in a bounded set

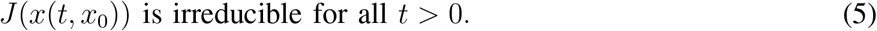

This assumption implies that the system (3) is strongly 2-cooperative.

Our main result shows that under these assumptions every trajectory of (3) that remains in a compact set has a well-ordered behavior. More precisely, it satisfies a Poincaré-Bendixson property.

### Theorem 1.

*Pick an initial condition* 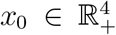 *and consider the omega limit set ω*(*x*_0_) *of the solu-tion x*(*t, x*_0_) *of* (3). *If ω*(*x*_0_) *is compact then the dynamics on ω*(*x*_0_) *is topologically conjugate to the dynamics of a compact invariant set of a Lipschitz-continuous vector field in* ℝ^2^.

### Example 1.

*Consider the system* (1) *described in the introduction, analyzed on the nonnegative orthant* 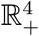*, which is a positively invariant set for its dynamics. Its Jacobian is*

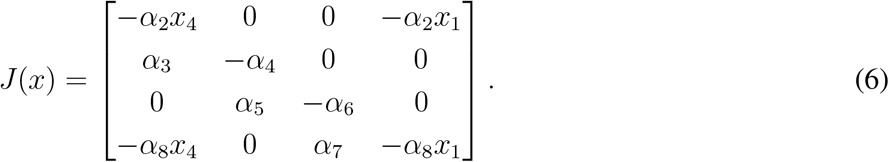

*Thus, it satisfies the assumed sign pattern*.

*We now show that* (5) *holds. Fix* 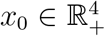*, and let x*(*t*) ≔ *x*(*t, x*_0_) *denote the corresponding solution of* (1)*. We assume that x*(*t, x*_0_) *remains in a bounded set, that is, there exists r* > 0 *such that*

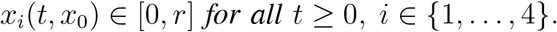

*From the structure of the matrix J in* (6)*, it follows that J* (*x*) *is irreducible provided that x*_1_ > 0*. We show that x*(*t*) *cannt be zero for any positive time, for any trajectory. More generally, suppose that* 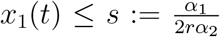 *for some t. Since x*_4_(*t*) ≤ *r for all t, it follows that* 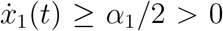*. This implies that there exist ε, T* > 0 *such that x*_1_(*t*) ≥ *εt for all t* ∈ [0, *T*] *and therefore x*_1_(*t*) ≥ *s for all t* ∈ [*T,* ∞). *Thus J* (*x*(*t, x*_0_)) *is indeed irreducible for all t* > 0.

*Clearly,* (1) *admits a unique equilibrium point at*

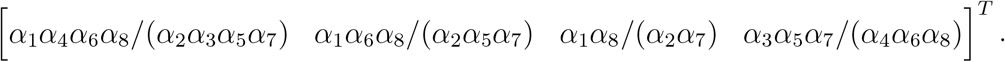

*Fig. 1 depicts the trajectory of* (1) *for the parameter values*

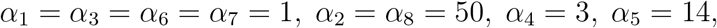

*and the initial condition x*_0_ = 0*. In this case the equilibrium is* [3/14 1/14 1 7/75]^*T*^. *It may be seen that the trajectory does not converge to the equilibrium and converges to a limit cycle.*

**Fig. 1.**
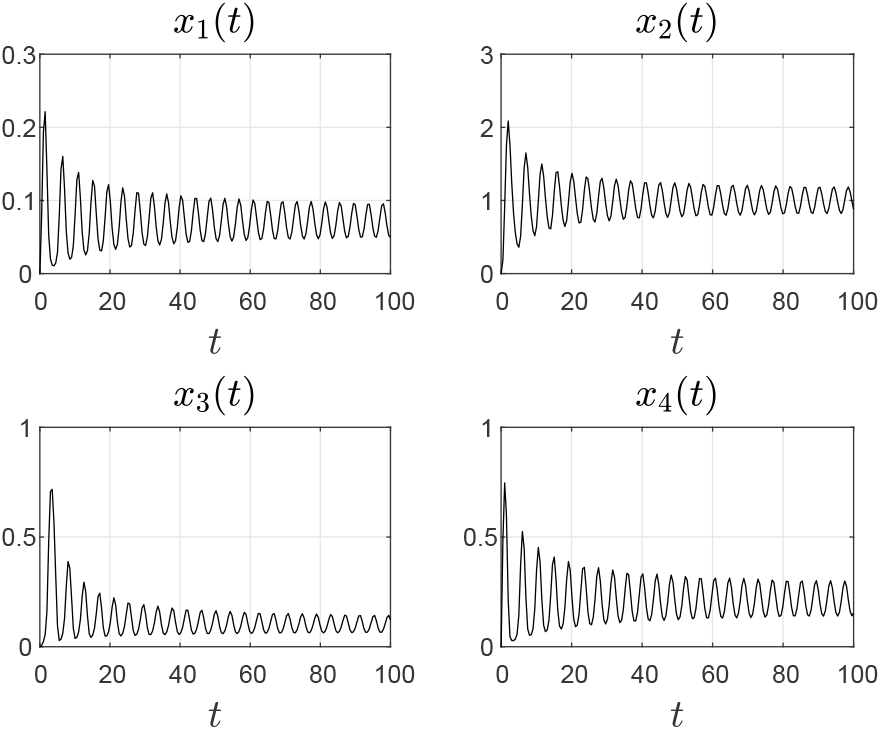
The state-variables of the system in Example 1.

The next section reviews some known definitions and results that are used in the proof of our main result.

## III. PRELIMINARIES

We begin by reviewing several known results on the number of sign variations in a vector, and linear mappings that preserve this number.

### A. Sign variations

For *y* ∈ ℝ^*n*^, let *s*^−^(*y*) denote the number of sign variations in *y* after deleting all its zero entries (with *s*^−^(0) defined as zero). For example, for *n* = 4 and *z* ≔ [1 0 0 −1]^T^, *s*^−^(*z*) = 1. Let *s*^+^(*y*) denote the maximal possible number of sign variations in *y* after replacing each zero entry with either 1 or −1. For example, for *n* = 4 and *z* ≔ [1 0 0 −1]^*T*^, *s*^+^(*z*) = 3. Note that these definitions imply that

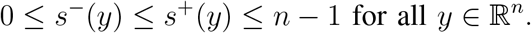

The following result provides information on the number of sign variations in the limit of a sequence of vectors.

#### Lemma 1.

*[21, Ch. 3] Let v*^*k*^, *k* = 1, 2, …, *be a sequence of vectors in* ℝ^*n*^. *If v* = lim_*k*→∞_*v*^*k*^ *then*

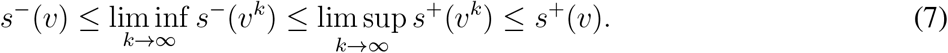

The proof follows from the fact that zero entries that appear in the limit vector *v* can only decrease [increase] *s*^−^ [*s*^+^].

We recall a useful duality relation between *s*^*−*^ and *s*^+^. Let

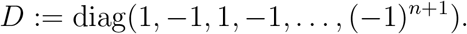

Then

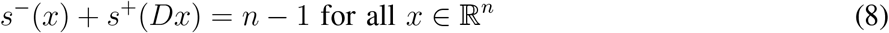

(see e.g. [21, Ch. 3]).

We can now define the sets of vectors with no more than *k*−1 sign variations. For any *k* ∈ {1, …, *n*−1}, let

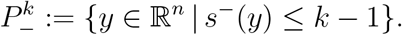

Then

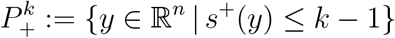

is the interior of 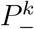 [29]. For example,

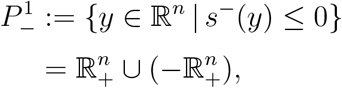

and

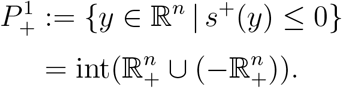

Note that this example shows in particular that the sets 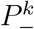 and 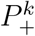 are in general *not* convex sets. For a geometric analysis of these sets, see [29].

Recall that a set *C* ⊆ ℝ^*n*^ is called a *cone of rank k* (see e.g. [13], [23]) if:

1. *C* is closed,
2. *x* ∈ *C* implies that *αx* ∈ *C* for all *α* ∈ ℝ, and
3. *C* contains a linear subspace of dimension *k* and no linear subspace of higher dimension.

For example, it is straightforward to see that 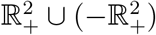 (and, more generally, 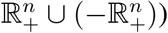) is a cone of rank 1. Thus, cooperative systems admit an invariant cone of rank 1.

A cone *C* of rank *k* is called *solid* if its interior is nonempty, and *k-solid* if there is a linear subspace 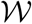 of dimension *k* such that 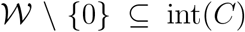. In the context of dynamical systems, such cones are important because trajectories of dynamical systems that are confined to *C* can be projected to the linear subspace *W* [23]. Roughly speaking, if this projection is one-to-one then the trajectories must satisfy the same properties as trajectories in a *k*-dimensional space.

It was shown in [29] that for any *k* ∈ {1, 2, …, *n* − 1} the set 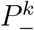 is a *k*-solid cone, and its complement

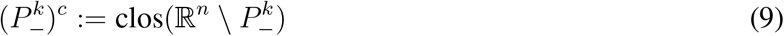

is an (*n* − *k*)-solid cone.

### B. Linear mappings that preserve the number of sign variations

A matrix is called *sign regular of order k* (*SR*_*k*_) if all its *k*-minors are either nonnegative or all are nonpositive. It is called *strictly sign regular of order k* (*SSR*_*k*_) if it is *SR*_*k*_ and all its *k* minors are nonzero (thus, they are either all positive or all negative). A matrix *A* ∈ ℝ^*n*×*m*^ is called *sign regular* (*SR*) if it is *SR*_*k*_ for all *k* ∈ {1, …, min{*n, m*}}, and *strictly sign regular* (*SSR*) if it is *SSR*_*k*_ for all *k* ∈ {1, …, min{*n, m*}}. These matrices have interesting sign variation diminishing properties (VDPs).

#### Theorem 2.

*[11] Let A* ∈ ℝ^*n*×*m*^ *be a matrix of full rank. Then A is SSR if and only if*

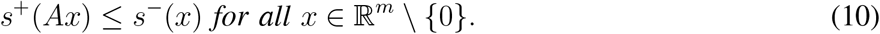

#### Example 2.

*Consider the case n* = *m* = 2, *that is,* 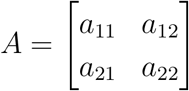. *Suppose that* (10) *holds. Taking x* = [1 0]^*T*^, *yields*

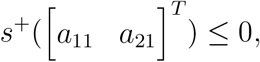

*so a*_11_, *a*_21_ *are either both positive or both negative. Similarly, Taking x* = [0 1]^*T*^ *implies that a*_12_, *a*_22_ *are either both positive or both negative. Now taking x* = [1 *c*]^*T*^, *with c* > 0, *implies that all the entries of A are either all positive or all negative, so A is SSR*_1_. *By the full rank assumption,* det(*A*) ≠ 0, *so A is also SSR*_2_, *and thus it is SSR.*

*Conversely, suppose that A is SSR. Then all its entries are either all positive or all negative. Seeking a contradiction, assume that* (10) *does not hold. Then there exists x* ≠ 0 *such that s*^+^(*Ax*) > *s*^−^(*x*), *that is, s*^+^(*Ax*) = 1 *and s*^−^(*x*) = 0*. We may assume that x*_1_, *x*_2_ ≥ 0. *Let y* ≔ *Ax. Then s*^+^(*y*) = 1 *implies that y*_1_*y*_2_ ≤ 0*. Since x*_1_, *x*_2_ ≥ 0 *and all the entries of A have the same strict sign, this implies that y* = 0*. Since A is SSR*_2_, *it is nonsingular, so x* = 0*. This contradiction implies that* (10) *holds.*

Note that if we drop the full rank assumption then Thm. 2 does not hold. For example, condition (10) holds for 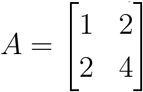, yet *A* is not *SSR*.

The most important examples of *SR* [*SSR*] matrices are the totally nonnegative [totally positive] matrices, i.e. matrices with all minors nonnegative [positive].

For our purposes, we require a more specific VDP.

#### Theorem 3.

*[5] Let A* ∈ ℝ^*n*×*n*^ *be a nonsingular matrix. Pick k* ∈ {0, …, *n* − 1}*. Then the following two conditions are equivalent:*

1. *For any x* ∈ ℝ^*n*^ \ {0} *with s*^−^(*x*) ≤ *k, we have*

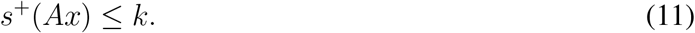
2. *A is SSR*_*k*+1_.

Note that condition (11) does not necessarily imply that *s*^+^(*Ax*) ≤ *s*^−^(*x*). Rather, it implies that *A* maps 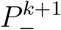 to 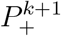.

The next subsection provides a tool for building bases of subspaces contained in 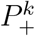, that is, in 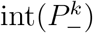.

### C. Oscillatory matrices

A matrix is called totally nonnegative (TN) if all its minors are nonnegative and totally positive (TP) if they are all positive. These matrices have many special and important properties [21], [8]. A matrix *A* ∈ ℝ^*n*×*n*^ is called *oscillatory* if it is TN and there exists an integer *k* ≥ 1 such that *A*^*k*^ is TP [11]. For example, if *A* is TP then it is oscillatory. Oscillatory matrices have a special spectral structure.

#### Theorem 4.

*[11], [22] If A* ∈ ℝ^*n*×*n*^ *is an oscillatory matrix then its eigenvalues are all real, positive, and distinct. Order the eigenvalues as λ*_1_ > *λ*_2_ > · · · > *λ*_*n*_ > 0, *and let u*^*k*^ ∈ ℝ^*n*^ *denote the eigenvector corresponding to λ*_*k*_. *For any* 1 ≤ *i* ≤ *j* ≤ *n, let* 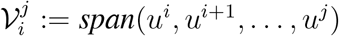*. Then*

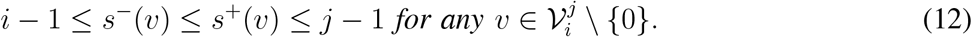

Thus, for any set in the form

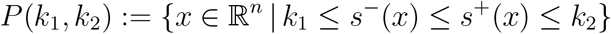

the eigenvectors of an oscillatory matrix can be used to provide an explicit representation of a sub-space 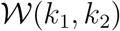 such that 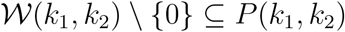.

Note that (12) implies in particular that

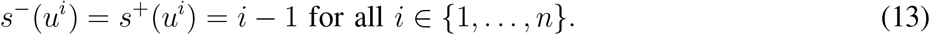

#### Example 3.

*Consider the n* × *n tridiagonal matrix*

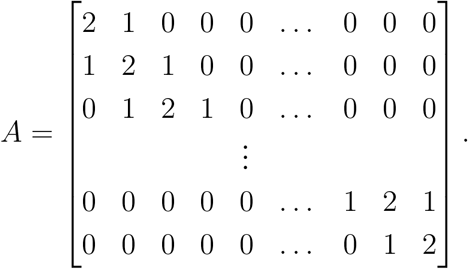

*It is well-known [8, Chapter 0] that A is oscillatory. Its eigenvalues λ*_*k*_ *and corresponding eigenvectors u*^*k*^, *k* = 1, …, *n, are given by*

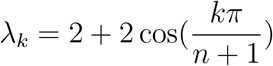

*and*

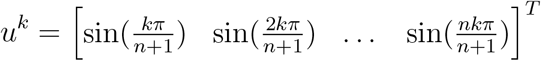

*(see, e.g. [30]). We conclude that the eigenvactors satisfy* (12)*. Note that since A is also symmetric, the u*^*i*^*s are orthogonal, so* 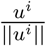, *i* = 1, …, *n, is an orthonormal set of vectors.*

*As a concrete example, take n* = 4*. Then the eigenvalues and eigenvectors are*

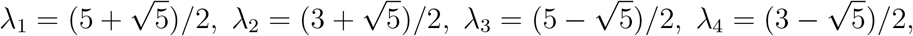

*and*

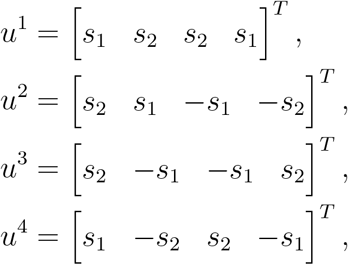

*with* 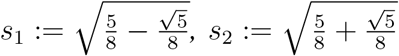, *and it is easy to verify that these vectors satisfy* (13).

The next result provides information on the number of sign changes of every non-zero vector in a subspace.

#### Proposition 1.

*[11, Ch. V] Let* 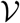 *be an m-dimensional subspace of* ℝ^*n*^, *with m* < *n. Then, the following statements are equivalent*:

1. *s*^+^(*v*) ≤ *m* − 1 *for all* 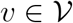;
2. *for every basis u*^1^, …, *u*^*m*^ *of* 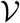, *the matrix U* ≔ [*u*^1^ *… u^m^*] ∈ ℝ^*n*×*m*^ *has all minors of order m nonzero and of the same sign.*

#### Example 4.

*Consider the osci tory matrix in Example 3 with n* = 3 *. Its first two eigenvectors are* 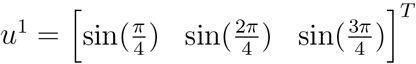 *and* 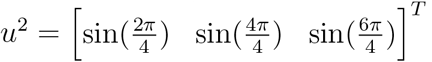 *. Thus,*

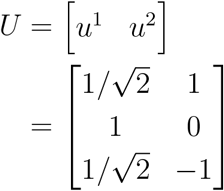

*and the minors of order* 2 *are* 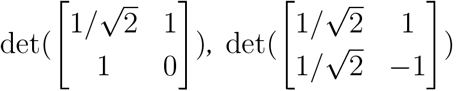, *and* 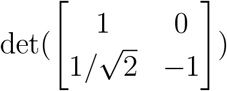. *These are all negative, so we conclude that for any c*_1_, *c*_2_ ∈ ℝ, *that are not both zero,*

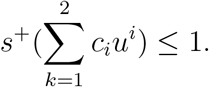

#### Remark 1.

*Combining Prop. 1 and* (8) *implies the following. Let* 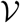 *be an m-dimensional subspace of ℝ*^*n*^, *with m < n. Then, the following statements are equivalent*:

1. *s*^+^(*v*) ≥ *n* − *m for all* 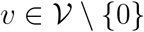;
2. *for every basis u*^1^, …, *u*^*m*^ *of* 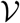, *the matrix D* [*u*^1^ … *u*^*m*^] ∈ ℝ^*n*×*m*^ *has all minors of order m nonzero and of the same sign*.

### D. Projections in cones that contain a subspace

The analysis of dynamical systems with invariant cones of rank *k* is based on projecting the flow of the *n*-dimensional original system to the flow of a *k*-dimensional system. The next result will be used to prove that the projection admits an inverse on a certain set.

By a *cone* in ℝ^*n*^ we mean a set *Q* ⊆ ℝ^*n*^ with the property that for all *r* > 0 and all *q* ∈ *Q*, *rq* ∈ *Q*. Let *Q*^*c*^ ≔ ℝ^*n*^\*Q*.

#### Lemma 2.

*Suppose that Q is an open cone and that there is a linear subspace* 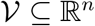 *such that* 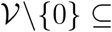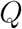*. Pick any complement* 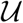 *of* 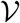, *meaning a linear subspace such that*

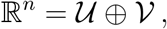

*and let* 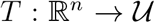 *be the projection on* 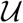 *along* 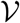*. In other words, given any x* ∈ ℝ^*n*^ *and its unique decomposition x* = *u* + *v with* 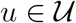 *and* 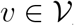, *T x* ≔ *u. Then, there is some k* > 0 *such that*

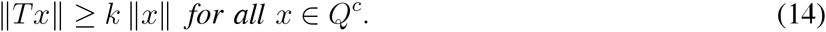

#### Remark 2.

*Note that this implies that if* Ω *is a set such that for any x, y* ∈ Ω *we have x* − *y* ∈ *Q*^*c*^ *then* ‖*T* (*x* − *y*)‖ ≥ *k* ‖*x* − *y*‖, *i.e. the restriction of T to* Ω *admits an inverse T* ^−1^ : *T* (Ω) → Ω *that is Lipschitz.*

Proof. Assume on the contrary that there are a sequence of elements *x*^*k*^ ∈ *Q*^*c*^ and positive numbers *ε*_*k*_ → 0 such that

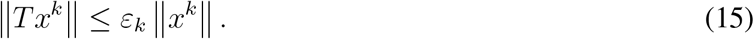

Let *x*^*k*^ = *u*^*k*^ + *v*^*k*^, 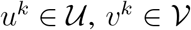. Then (15) gives ‖*u*^*k*^‖ ≤ *ε*_*k*_‖*x*^*k*^‖ for all *k*. Introduce

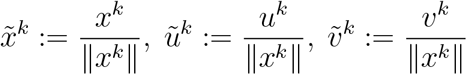

and note that 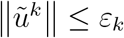, so 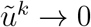 as *k* → ∞.

On the other hand, since every 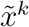 has unit norm, we may assume, taking a subsequence if necessary, that 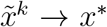 as *k* → ∞, for some *x*^*^ ∈ ℝ^*n*^ of unit norm. Since 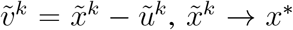, and 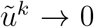, it follows that 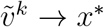 as *k* →∞. Since 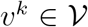 and 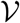 is a subspace also 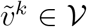 for all *k*. Moreover, as 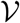 is closed (being a subspace) and *x*\* ≠ 0 (it has unit norm) it follows that 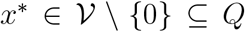. Since *Q* is open and 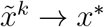, it follows that there is some *k* such that 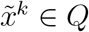. Since *Q* is a cone, also 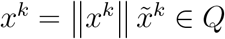. This co1ntra1dicts the assumption that *x*^*k*^ ∈ *Q*^*c*^, and thus completes the proof. ◻

#### Example 5.

*Let n* = 2, *Q* = {*x* ∈ ℝ^2^ | |*x*_2_| > |*x*_1_|}, 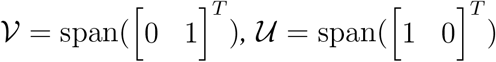, *and let* ‖·‖ *denote the Euclidean norm. Here T* ([*x*_1_ *x*_2_]^*T*^) = *x*_1_, *and Q*^*c*^ = {*x* ∈ ℝ^2^ | |*x*_2_ | ≤ |*x*_1_ |}*. Then Lemma 2 implies that* (14) *holds for some k* > 0*. Indeed, it is easy to verify directly that* (14) *holds for* 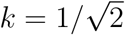.

Lemma 2 does not require *T* to be an orthogonal projection. The next example demonstrates this.

#### Example 6.

*Let*

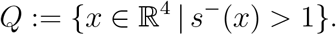

*This is clearly an open cone (note that it is the complement of the closed set* 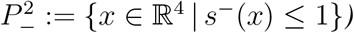. *Let* 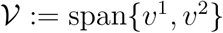, *where*

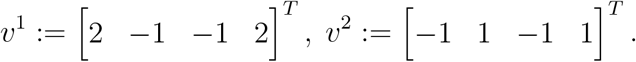

*Using Remark 1 it is straightforward to verify that* 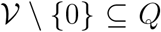*. A possible complement of* 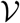 *is* 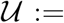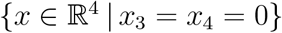*. The set*

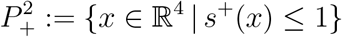

*satisfies* 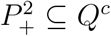, *so* (14) *implies that there exists k* > 0 *such that for any x* ∈ ℝ^4^, *with s*^+^(*x*) ≤ 1, *we have*

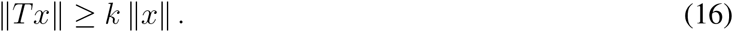

*As a specific example, take x* = [*x*_1_ 0 *x*_3_ *x*_4_]^T^, *with x* < 0, *and x*_3_, *x*_4_ > 0*. Then* 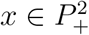 *. Its unique decomposition as x* = *u* + *v, with* 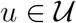 *and* 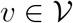, *is*

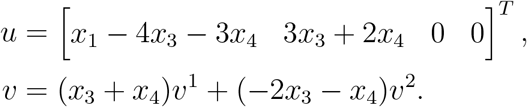

*Thus, in this case* (16) *implies in particular that there exists k* > 0 *such that*

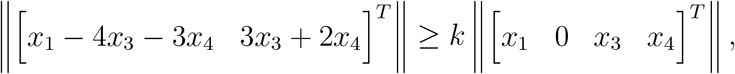

*for all x*_1_ < 0, *and x*_3_, *x*_4_ > 0.

The next section provides the proof of Thm. 1. This is based on several auxiliary results that may be of independent interest.

## IV. PROOF OF MAIN RESULT

Our first auxiliary result shows that the vector of derivatives 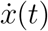 of (3) satisfies a special sign pattern property.

### Lemma 3.

*Pick an initial condition* 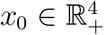 *and consider the solution x*(*t*) ≔ *x*(*t, x*_0_) *of* (3)*. Suppose that* 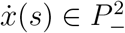 *for some s* ≥ 0*. Then* 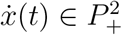 *for all t > s.*

### Example 7.

*Consider* (1) *with α*_*i*_ = 1 *for all i. Let x*_0_ ≔ [2 1 2.2 1]^*T*^. *For this initial value x*(*t, x*_0_) *converges to the equilibrium* [1 1 1 1]^*T*^. *Fig. 2 depicts* 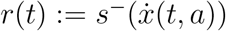 *as a function of t* ∈ [0, 12]. *It may be seen that r*(0) = 3, *then drops to r*(*t*) = 1 *at t* ≈ 0.8936*. From here on, the value of r*(*t*) *changes values between* 0 *and* 1. *This agrees with the fact that if* 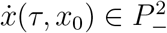 *for some τ then* 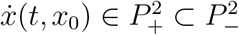 *for all t > τ*.

*Proof of Lemma 3:* Let *x*(*t*) = *x*(*t, x*_0_) be a solution of (3). The associated variational system along this solution is:

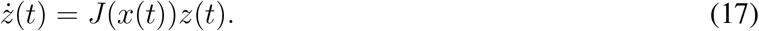

Thus,

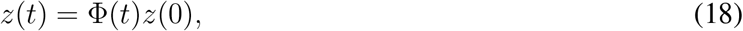

where

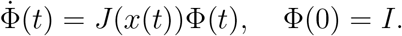

Let (Φ(*t*))^(2)^ denote the matrix that includes all the 2 × 2 minors of Φ(*t*) ordered in a lexicographic order. This has dimensions 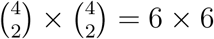. Then (Φ(*t*))^(2)^ satisfies the differential equation:

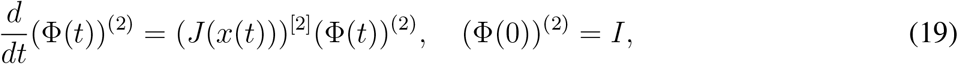

where *B*^[2]^ denotes the 2nd additive compound of the matrix *B* (see, e.g. [24], [16]). Recall (see e.g. [24]) that for *A* ∈ ℝ^4×4^ the 2nd additive compound is

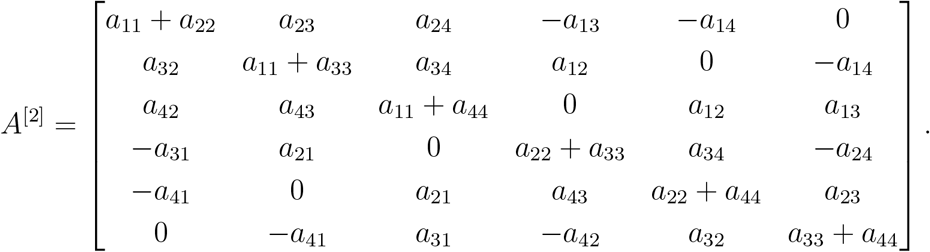

Combining this with (4) gives

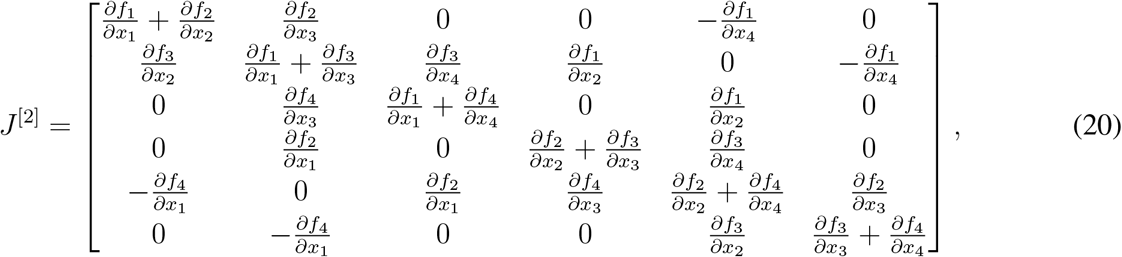

and our assumptions on the sign pattern of the Jacobian imply that (*J* (*x*(*t*)))^[2]^ is Metzler for all *t* ≥ 0. Thus (19) implies that

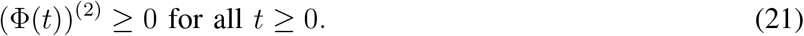

In other words, all 2 × 2 minors of Φ(*t*) are non-negative for all *t* ≥ 0. Furthermore, (5) implies that all 2 × 2 minors of Φ(*t*) are positive for all *t* > 0.

By Thm. 3, if all the 2 × 2 minors of a matrix *B* are positive then for any x ≠ 0 with *s*^−^(*x*) ≤ 1, we have *s*^+^(*Bx*) ≤ 1. Applying this to (18) and using the fact that 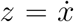 satisfies the variational equation completes the proof.

**Fig. 2.**
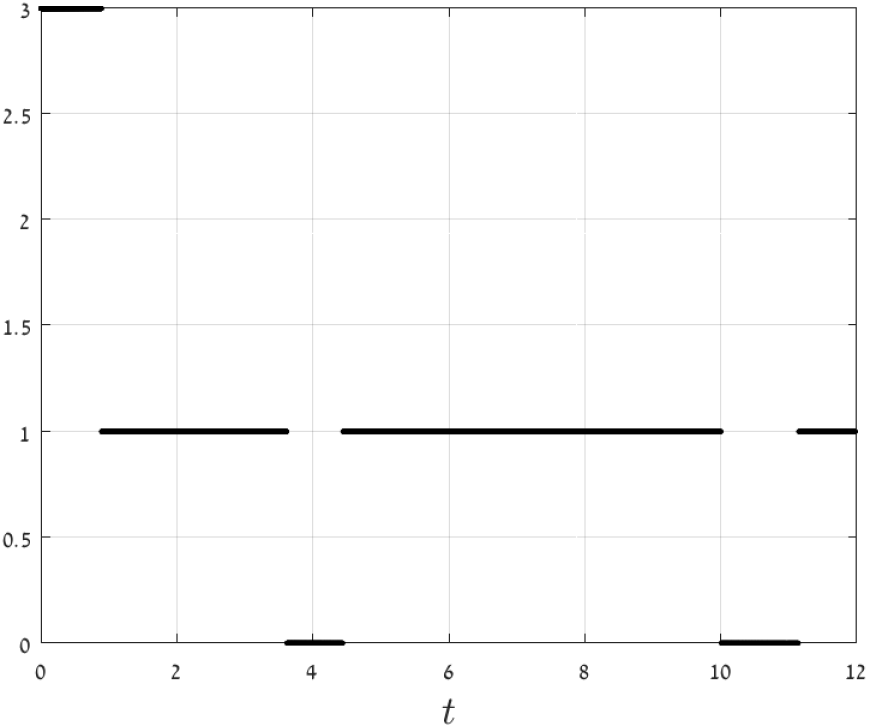
The function 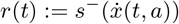 as a function of *t* in Example 7.

### Remark 3.

*Our goal in computing J*^[2]^ *was to prove that* (19) *is a cooperative dynamical system. We note in passing that under certain conditions J* ^[2]^ *can also be used to rule out the existence of limit cycles [18]. This is based on a contraction argument (see e.g. [3]). For another application of the 2nd additive compound, see [14].*

Let *D* ≔ diag(1, −1, 1, −1). Fix 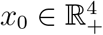 so that *x*(*t, x*_0_) is not a constant solution. Note that 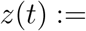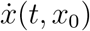 satisfies the variational equation. Lemma 3 implies that we can classify the trajectory *x*(*t, x*_0_) into one of two types:

Type 1: If 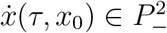 for some *τ* ≥ 0 then 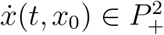 for all *t > τ*;
Type 2: If 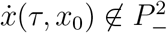 for all *τ* ≥ 0 then 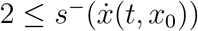 for all *t* ≥ 0, and using the duality

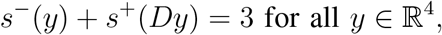

implies that 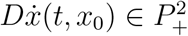 for all *t* ≥ 0.

Our next goal is to transform these properties of 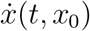 to corresponding properties of *x*(*t*_2_, *x*_0_) − *x*(*t*_1_, *x*_0_) with *t*_2_ *> t*_1_ ≥ 0. Following [23], we first do this for solutions that are closed orbits.

### Lemma 4.

*Let* γ *be a closed orbit corresponding to a periodic solution x*(*t* + *T, x*_0_) = *x*(*t, x*_0_) *for all t* ≥ 0, *where T* > 0 *is the minimal period. If x*(*t, x*_0_) *is Type 1 then*

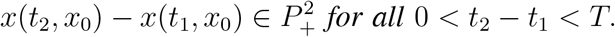

*If x*(*t, x*_0_) *is Type 2 then*

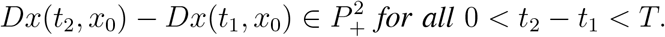

*Proof of Lemma 4:* Suppose that *x*(*t, x*_0_) is Type 1. Seeking a contradiction, assume that there exist two distinct points *p, q* ∈ γ such that 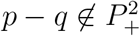. Let *τ*_1_, *τ*_2_ be such that 0 < *τ*_2_ − *τ*_1_ *< T*, *x*(*τ*_1_, *x*_0_) = *p* and *x*(*τ*_2_, *x*_0_) = *q*. Note that by adding *kT* to *τ*_1_, *τ*_2_ we may assume that *τ*_1_, *τ*_2_ are arbitrarily large. Since *x*(*t, x*_0_) is Type 1, 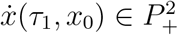, and Lemma 1 implies that

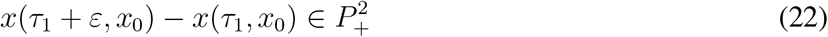

for all *ε* > 0 sufficiently small. Hence, we may actually assume that

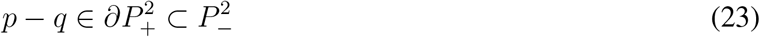

and since 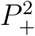 is an open set

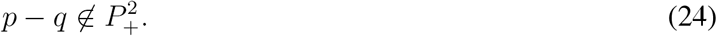

Let *z*(*t*) ≔ *x*(*t, p*) − *x*(*t, q*). Then

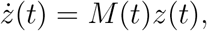

with 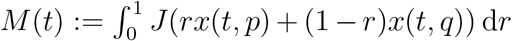. Note that *M* (*t*) satisfies the same sign pattern as *J* does. Thus, if 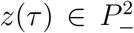 for some *τ* ≥ 0 then 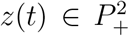 for all *t > τ*. Eq. (23) implies that 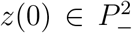, so 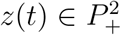 for all *t* > 0 and in particular 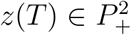. Thus, 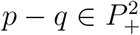. This contradiction completes the proof in the case where *x*(*t, x*_0_) is Type 1. The proof in the case where *x*(*t, x*_0_) is Type 2 is similar.

An interesting implication of Lemma 4 is that there exists a one-to-one projection of closed orbits of the four-dimensional system to a two-dimensional subspace.

### Lemma 5.

*There exists a two-dimensional subspace* 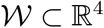 *such that* 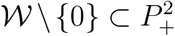. *Let* γ *be a closed orbit corresponding to a periodic solution x*(*t* + *T, x*_0_) = *x*(*t, x*_0_) *for all t* ≥ 0, *where T* > 0 *is the minimal period. If x*(*t, x*_0_) *is Type 1 then the orthogonal projection of x*(*t, x*_0_), *t* ∈ [0, *T*), *to* 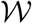 *is one-to one. If x*(*t, x*_0_) *is Type 2 then the orthogonal projection of Dx*(*t, x*_0_), *t* ∈ [0, *T*), *to* 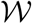 *is one-to one.*

### Example 8.

*Let* 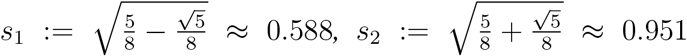 *. One possible choice for the subspace* 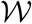, *that is based on the special spectral properties of oscillatory matrices (see Section III), is:*

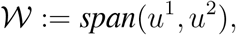

*where*

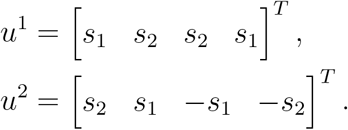

*Furthermore,* 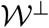 *,the orthogonal complement of* 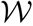, *satisfies*

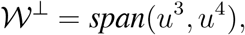

*with*

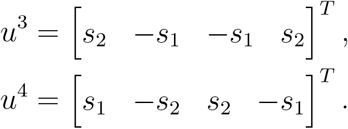

*Note that* (*u*^*i*^)^*T*^*u*^*j*^ = 0 *for all i* ≠ *j.*

*Proof of Lemma 5:* Suppose that *x*(*t, x*_0_) corresponds to a closed trajectory γ. Consider the case where *x*(*t, x*_0_) is Type 1. Seeking a contradiction, assume that there exist *p, q* ∈ γ, *with p* ≠ *q*, such that

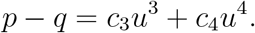

Then 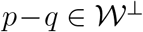, so *s*^+^(*p*−*q*) > 1 and this is a contradiction of Lemma 4. We conclude that the orotognal projection of γ onto the 2-dimensional subspace 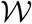 is one-to-one. The proof in the case where *x*(*t, x*_0_) is Type 2 is similar.

### Remark 4.

*Let e*^*i*^, *i* = 1, …, 4, *denote the ith canonical vector in* ℝ^4^*. Consider the subspace spanned by two canonical vectors, say,* 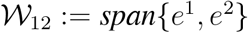*. Then clearly* 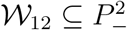, *but* 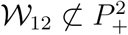. *The orthogonal projection to* 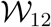 *is T*_12_(*x*) ≔ [*x*_1_ *x*_2_]^*T*^, *i.e. T* (*x*(*t*)) *amounts to taking the first two-state-variables. The proof of Lemma 5 remains valid if we replace T, the orthogonal projection to* 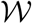, *by T*_12_*. Indeed, consider for example the case where x*(*t, x*_0_) *is Type* 1*. Assume that there exist p, q* ∈ γ, *with p ≠ q, such that*

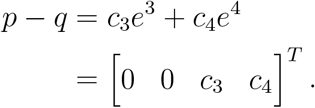

*Then* 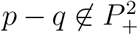, *and this contradiction shows that T*_12_(γ) *is one-to-one. As an example, consider again the periodic solution described in Example 1. Fig. 3 depicts the trajectory x*_1_(*t*), *x*_2_(*t*) *(i.e. the orthogonal projection of x*(*t, a*) *onto T*_12_*), for x*_0_ = 0*. It may be seen that the curve does not intersect itself, and this agrees with the fact that the projection is one-to-one.*

*However, in what follows we will be interested in projecting the omega limit set of a solution x*(*t, x*_0_) *that is not necessarily periodic to a two-dimensional subspace, and in this case we can prove that the projection T is one-to-one, but we cannot prove the same for T*_12_.

**Fig. 3.**
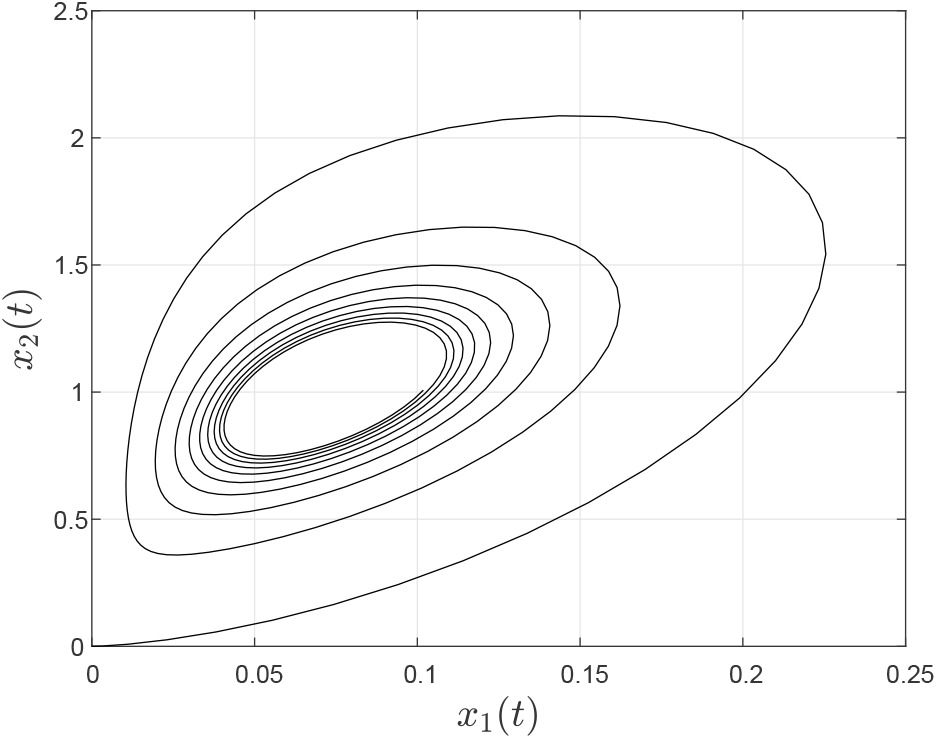
The curve (*x*_1_(*t*), *x*_2_(*t*)) described in Remark 4.

The next step is to analyze trajectories that are not necessarily closed orbits.

### Lemma 6.

*Pick* 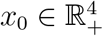 *such that x*(*t, x*_0_) *remains in a bounded set. Let ω*(*x*_0_) *denote the omega limit set of x*_0_*. Then at least one of the following two alternatives holds.*

a. *any p, q* ∈ *ω*(*x*_0_) *satisfy* 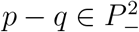;
b. *any p, q* ∈ *ω*(*x*_0_) *satisfy* 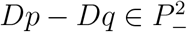.

*Proof:* If *x*(*t, x*_0_) is Type 1 then [23, Theorem 2] implies (a). If *x*(*t, x*_0_) is Type 2 then 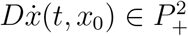 for all *t* ≥ 0. Since *D*^−1^ = *D*, this implies that

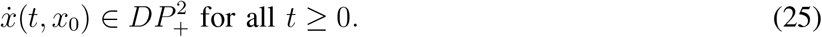

It is straightforward to verify that 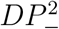 is a cone of rank 2, and clearly (25) trivially implies that

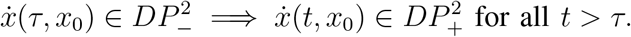

Thus, we can apply the results of Sanchez to the dynamical system and the cone 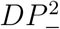 of rank 2 to conclude that 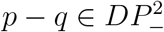.

We can now prove our main result.

*Proof of Thm. 1:* Pick 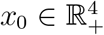 such that *x*(*t, x*_0_) remains in a bounded set. Then the omega limit set *ω* = *ω*(*x*_0_) is non-empty and compact.

Consider the case where *x*(*t, x*_0_) is Type 1. Let *T* denote the orthogonal projection operator onto 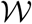. Assume that there exist *p, q* ∈ *ω*, with *p* ≠ *q*, such that

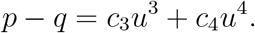

Then 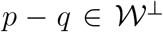, so *s*^−^(*p* − *q*) > 1 and this contradicts Lemma 6. We conclude that the projection of *ω* onto the 2-dimensional subspace 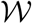 is one-to-one. Thus, *T*_*ω*_, the restriction of *T* to *ω*, is a Lipschitz homeomorphism of *ω* onto a compact subset of 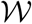.

Applying Lemma 2 with *Q* ≔ {*x* ∈ ℝ^4^ | *s*^−^(*x*) > 1}, (so 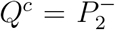), 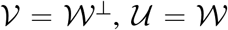 implies that there exists *k* > 0 such that

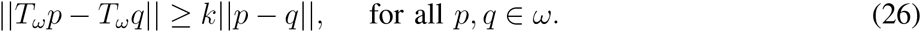

Therefore, 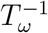 exists and is Lipschitz on *T* (*ω*).

Pick *p* ∈ *ω*. Consider the vector field

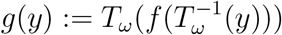

defined on *T* (*ω*), where *f* is the vector field of (3). The vector field *g* can be extended to a Lipschitz vector field on ℝ^2^ [17]. The solution of 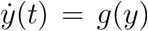, with *y*(0) = *T*_*ω*_ (*p*), is *y*(*t*) = *T*_*ω*_ (*x*(*t, p*)). Note that *T* (*ω*) is a compact invariant set for the *y* system.

We conclude that the flow on a compact omega limit set of (3) is topologically equivalent to the flow on a compact invariant set of a Lipschitz system of differential equations in ℝ^2^. This completes the proof when *x*(*t, x*_0_) is Type 1.

The proof when *x*(*t, x*_0_) is Type 2 is similar.

## V. CONCLUSION

We applied the recently developed theory of *k*-cooperative dynamical systems to analyze an important model from systems biology. We showed that the model is a 2-cooperative dynamical system and used this to infer a Poincaré-Bendixson property for every solution that remains in a compact set. Note that in general the results for cooperative systems are of the form “for almost any initial condition some property holds”. In our case, the special structure of four-dimensional 2-cooperative systems yields a stronger result.

The analysis using the theory of *k*-cooperative dynamical systems has several advantages. First, just like in cooperative systems, the condition for *k*-cooperativity is a *sign pattern* condition on the Jacobian [29]. This is useful in fields like systems biology where one often knows if a reaction is either inhibitory or excitatory. Second, the omega limit set of an *n*-dimensional *k*-cooperative dynamical system can be projected to that of a *k*-dimensional system, and the projection is explicit. An intersting topic for further research is to study the implications of this explicit proejction.

## REFERENCES

[1] D. K. Agrawal, R. Marshall, M. A. Al-Radhawi, V. Noireaux, and E. D. Sontag, “Some remarks on robust gene regulation in a biomolecular integral controller,” in Proc. 2019 IEEE Conf. Decision and Control, 2019, to appear.

[2] D. K. Agrawal, R. Marshall, V. Noireaux, and E. D. Sontag, “In vitro implementation of robust gene regulation in a synthetic biomolecular integral controller,” Nature Communications, 2019, in press.

[3] Z. Aminzare and E. D. Sontag, “Contraction methods for nonlinear systems: A brief introduction and some open problems,” in Proc. 53rd IEEE Conf. on Decision and Control, Los Angeles, CA, 2014, pp. 3835–3847.

[4] S. K. Aoki, G. Lillacci, A. Gupta, A. Baumschlager, D. Schweingruber, and M. Khammash, “A universal biomolecular integral feedback controller for robust perfect adaptation,” Nature, vol. 570, pp. 533–537, 2019.

[5] T. Ben-Avraham, G. Sharon, Y. Zarai, and M. Margaliot, “Dynamical systems with a cyclic sign variation diminishing property,” IEEE Trans. Automat. Control, 2019, to appear. [Online]. Available: https://ieeexplore.ieee.org/document/8706539

[6] C. Briat, A. Gupta, and M. Khammash, “Antithetic integral feedback ensures robust perfect adaptation in noisy biomolecular networks,” Cell Syst., vol. 2, pp. 15–26, 2016.

[7] A. S. Elkhader, “A result on a feedback system of ordinary differential equations,” J. Dynam. Diff. Equations, vol. 4, pp. 399–418, 1992.

[8] S. M. Fallat and C. R. Johnson, Totally Nonnegative Matrices. Princeton, NJ: Princeton University Press, 2011.

[9] L. Farina and S. Rinaldi, Positive Linear Systems: Theory and Applications. John Wiley, 2000.

[10] L. Feng, Y. Wang, and J. Wu, “Semiflows “monotone with respect to high-rank cones” on a Banach space,” SIAM J. Math. Anal., vol. 49, no. 1, pp. 142–161, 2017.

[11] F. R. Gantmacher and M. G. Krein, Oscillation Matrices and Kernels and Small Vibrations of Mechanical Systems. Providence, RI: American Mathematical Society, 2002, translation based on the 1941 Russian original.

[12] J. Huang, A. Isidori, L. Marconi, M. Mischiati, E. D. Sontag, and W. M. Wonham, “Internal models in control, biology and neuroscience,” in Proc. 2018 IEEE Conf. Decision and Control, 2018, pp. 5370–5390.

[13] M. A. Krasnoselskii, E. A. Lifshitz, and A. V. Sobolev, Positive Linear Systems: The Method of Positive Operators. Berlin: Heldermann Verlag, 1989.

[14] M. Y. Li and L. Wang, “A criterion for stability of matrices,” J. Math. Anal. Appl., vol. 225, pp. 249–264, 1998.

[15] J. Mallet-Paret and H. L. Smith, “The Poincaré-Bendixson theorem for monotone cyclic feedback systems,” J. Dyn. Differ. Equ., vol. 2, no. 4, pp. 367–421, 1990.

[16] M. Margaliot and E. D. Sontag, “Revisiting totally positive differential systems: A tutorial and new results,” Automatica, vol. 101, pp. 1–14, 2019.

[17] E. J. McShane, “Extension of range of functions,” Bull. Amer. Math. Soc., vol. 40, pp. 837–842, 1934.

[18] J. S. Muldowney, “Compound matrices and ordinary differential equations,” The Rocky Mountain J. Math., vol. 20, no. 4, pp. 857–872, 1990.

[19] N. Olsman and F. Forni, “Antithetic integral feedback for the robust control of monostable and oscillatory biomolecular circuits,” 2019. [Online]. Available: https://arxiv.org/abs/1911.05732

[20] S. Pigolotti, S. Krishna, and M. H. Jensen, “Oscillation patterns in negative feedback loops,” Proceedings of the National Academy of Sciences, vol. 104, pp. 6533–6537, 2007.

[21] A. Pinkus, Totally Positive Matrices. Cambridge, UK: Cambridge University Press, 2010.

[22] A. Pinkus, “Spectral properties of totally positive kernels and matrices,” in Total Positivity and its Applications, M. Gasca and C. A. Micchelli, Eds. Dordrecht: Springer Netherlands, 1996, pp. 477–511.

[23] L. A. Sanchez, “Cones of rank 2 and the Poincaré-Bendixson property for a new class of monotone systems,” J. Diff. Eqns., vol. 246, no. 5, pp. 1978–1990, 2009.

[24] B. Schwarz, “Totally positive differential systems,” Pacific J. Math., vol. 32, no. 1, pp. 203–229, 1970.

[25] J. Smillie, “Competitive and cooperative tridiagonal systems of differential equations,” SIAM J. Math. Anal., vol. 15, pp. 530–534, 1984.

[26] H. L. Smith, “Periodic tridiagonal competitive and cooperative systems of differential equations,” SIAM J. Math. Anal., vol. 22, no. 4, pp. 1102–1109, 1991.

[27] H. L. Smith, Monotone Dynamical Systems: An Introduction to the Theory of Competitive and Cooperative Systems, ser. Mathematical Surveys and Monographs. Providence, RI: Amer. Math. Soc., 1995, vol. 41.

[28] R. A. Smith, “The Poincaré-Bendixson theorem for certain differential equations of higher order,” Proc. Royal Society of Edinburgh: Section A Mathematics, vol. 83, no. 1-2, pp. 63–79, 1979.

[29] E. Weiss and M. Margaliot, “A generalization of linear positive systems with applications to nonlinear systems: Invariant sets and the Poincaré-Bendixson property,” 2019, submitted. [Online]. Available: https://arxiv.org/abs/1902.01630

[30] W.-C. Yueh, “Eigenvalues of several tridiagonal matrices,” Applied Mathematics E-Notes, vol. 5, pp. 66–74, 2005.

